# Modelling glioblastoma resistance to temozolomide. Combination of spheroid and mathematical models to simulate cellular adaptation in vitro

**DOI:** 10.1101/2023.11.24.568421

**Authors:** Marina Pérez-Aliacar, Jacobo Ayensa-Jiménez, Teodora Ranđelović, Ignacio Ochoa, Manuel Doblaré

## Abstract

Drug resistance is one of the biggest challenges in the fight against cancer. In particular, in the case of glioblastoma, the most lethal brain tumour, resistance to temozolomide (the standard of care drug for chemotherapy in this tumour), is one of the main reasons behind treatment failure and hence responsible for the poor prognosis of patients diagnosed with this disease.

In this paper, we combine the power of three-dimensional in vitro experiments of treated glioblastoma spheroids with mathematical models of tumour evolution and adaptation. We use a novel approach based on internal variables for modelling the acquisition of resistance to temozolomide that is observed in a group of treated spheroids in the experiments. These internal variables describe the cell’s phenotypic state, which depends on the history of drug exposure and affects cell behaviour. We use model selection to determine the most parsimonious model and calibrate it to reproduce the experimental data, obtaining a high level of agreement between the in vitro and in silico outcomes. A sensitivity analysis is carried out to investigate the impact of each model parameter in the predictions. More importantly, we show the utility of our model for answering biological questions, such as what is the intrinsic adaptation mechanism, or for separating the sensitive and resistant populations. We conclude that the proposed in silico framework, in combination with experiments, can be useful to improve our understanding of the mechanisms behind drug resistance in glioblastoma and to eventually set some guidelines for the design of new treatment schemes.

## 1 Introduction

Glioblastoma (GBM) is one of the most challenging diseases to treat in oncology [1]. Despite being rare (incidence of 3 cases per 100,000 person-years [2]), it is the most common malignant primary central nervous system tumour, accounting for 48.6% of them [3]. It is also the most lethal brain tumour, with a 5-year survival rate of 6.8% [4] and a median survival of around 14 months after diagnosis in patients having received the current standard treatment [5, 6].

This current treatment is the so-called Stupp protocol [7], which consists of maximal safe resection by surgery followed by radiotherapy with concomitant and adjuvant chemotherapy with temozolomide (TMZ), the only approved drug for this protocol. The TMZ schedule according to this protocol comprises six cycles of 28 days, in each of which TMZ is administered the first five days at a daily dose of 150-200 mg*/*m^2^. However, this treatment increases the median overall survival only from 12.1 to 14.3 months [7]. Since its approval in 2005, and despite considerable efforts, there have been no significant advances in the treatment or the survival rate of this tumour [8].

TMZ is an oral alkylating agent which acts on GBM cells by inducing epigenetic changes in them [9]. Epigenetics refers to the heritable changes in gene expression of cells that are not caused by alterations in the DNA sequence, hence leading to phenotypic plasticity [10]. Both epigenetic reprogramming and phenotypic plasticity have proven to be crucial in every step of cancer progression, to the point that in 2022 they were included among the cancer hallmarks [11]. In particular, the mechanism of action of TMZ consists in inducing methylation in several DNA bases. It affects proliferating cells inducing cell cycle arrest and hence, preventing cells from duplicating (that is, causing a cytostatic effect), eventually leading to apoptosis [12, 13]. However, more than 50% of the GBM patients do not respond to the treatment with TMZ [14] due to the intrinsic and acquired resistance of GBM to chemotherapy. Indeed, GBM cells are able to reverse the epigenetic changes induced by TMZ and recover their normal proliferative behaviour [15]. This induced resistance is predominantly associated with an over expression of O^6^-methylguanine-DNA methyltransferase (MGMT) protein, which can repair the DNA damage induced by TMZ, removing the methyl groups attached to the DNA [9, 16]. There are also other mechanisms of acquired resistance, such as the emergence of populations of cancer stem cells [17]. Chemotherapy has proven to drive cells towards stem-like phenotypes, associated with higher drug resistance [18].

To overcome the stagnation in the development in new treatments for this tumour, it is paramount to better understand the behaviour of GBM and the mechanisms that trigger chemoresistance. Three-dimensional culture models are the most appropriate tools to in vitro reproduce tumour behaviour and drug response since they are able to closely mimic the in vivo physiological conditions, both structural and functional [19]. Among them, spheroids are the most frequently used technique [4] due to the fact that they are easy to culture, and they are able to reproduce cell-cell interactions and the gradients present in solid tumours [20]. In particular, in this paper we focus on the results obtained by Ranđelović et al. in a previous work [21], where spheroids of the GBM commercial line U87 where treated with TMZ following the clinical schedule. In response to treatment, there were two different responses. While a group of treated spheroids exhibited a stagnation in their growth maintained throughout the whole experiment, another group developed resistance and was able to resume growing, despite TMZ administration.

In vitro experiments, such as the one described above, still present some limitations, specially in terms of time and cost. Mathematical models, although unable to completely replace in vitro experimentation, represent a great complement, providing further explanatory capacity and allowing to test new hypotheses in a cheap and quick way. In this line, the main aim of this work is to develop a mathematical model incorporating the acquisition of resistance in GBM, and to quantitatively reproduce the experimental results obtained in [21]. The model must take into account cellular adaptation to the microenvironment, to consider how epigenetic changes induced by TMZ result in the cytostatic effect, as well as the eventual repair of these changes leading to the development of resistance. The majority of existent drug resistance models tackle this issue with a multiphenotypic approach, in which a discrete number of phenotypes (usually sensitive and resistant cells) behave differently and transitions between them are defined [22, 23, 24, 25].Yet this kind of models fails to recreate the complex reality, in which cells go through a wide spectrum of intermediate phenotypes in their way to becoming resistant [26]. Contrary to this first approach, some models have been proposed in which phenotype or cell state is considered as a continuum field, such as in [26], where there is an extra dimension added to model the possible phenotypic states. In a previous work, we proposed a general framework to model cellular adaptation including cell phenotype as defined by a set of internal variables that describe cell state [27]. An evolution equation is derived for each internal variable, allowing an easy coupling with the microenvironment as well as with cell evolution.

Hereafter, we particularise this model to the problem of GBM adaptation to TMZ and calibrate its parameters to match the experimental data. However, when working with mathematical models, it becomes paramount to search for parsimony to avoid generating overly-complicated models that are not useful for drawing conclusions not derived directly from the initial hypotheses. For this purpose, in the calibration process, we apply a methodology for model selection among different candidates, based on the Bayesian Information Criterion (BIC) [28, 29]. Subsequently, we perform a local sensitivity analysis to test the robustness of the model to the fitting. Once this validated, we can interrogate the model about biological questions, such as the cause of the differences between the two populations of spheroids (those who develop resistance and those who do not) and the ways of separating both groups.

The structure of the paper is as follows. Section 2 describes the experimental data on which the paper is based and the mathematical model particularised for the problem in hands. It also dwells into the methodology followed to perform model selection, parameter fitting and the posterior sensitivity analysis. Then, the results are presented in Section 3. In the first place, the best model candidate and the resulting parameter fitting is detailed. Next, the sensitivity analysis results are presented and finally, we highlight the potential of models such as the one presented by addressing some questions of biological relevance. Section 4 discusses the implications of the results and the limitations of the framework. The paper closes with the main conclusions in Section 5.

## 2 Materials and methods

### 2.1 Experimental data

GBM spheroids of the cell line U-87 MG (U87) were grown by the non-adherent surface method, with an initial number of 1000 cells per spheroid. The spheroids were kept in suspension in 96 well plates with growth medium [21].

Spheroids in the control group were not treated with TMZ. Treated spheroids were subjected to two cycles of the clinical treatment scheme, i.e. 5-day treatment (with a daily dose of TMZ) followed by 23 days of rest. Hence, the evolution of the spheroids was followed during 56 days. The TMZ dose used in the experiments was 100 *μ*M, which is the maximum concentration measured in patient’s plasma [30].

To analyse the data, bright field images were acquired using a Nikon Eclipse Ti-E C1 confocal microscope and analysed using Fiji’s plugin SpheroidJ to measure the spheroid’s area [31].

The experimental results are presented in Figure 1. It can be seen that control spheroids grow throughout the whole experiment. Regarding treated spheroids, they have been divided in two groups according to their behaviour. The first group, baptised as TMZ-sensitive (TMZ-S) respond to treatment and stop growing around the fifth day, maintaining their size for the rest of experiment. The fact that the growth stops after day 5, while TMZ was administered since day 1 suggests the existence of a certain lag between the administration on the drug and its effect on cells. This can be due to the time that it takes for the compound to reach the nucleus and then induce cell cycle arrest. We can observe the cytostatic effect of TMZ, which is maintained over time. Conversely, the second group, named TMZ-resistant (TMZ-R), despite having the same behaviour than the TMZ-S group until day 21, from this day they develop resistance to TMZ and reactivate their growth. The development of resistance was confirmed through transcriptomic analysis. For further details on the experimental methods and results, the reader is referred to [21, 32]. Figure 1 summarises the temporal evolution of the different groups of spheroids in the experiment.

**Figure 1:**
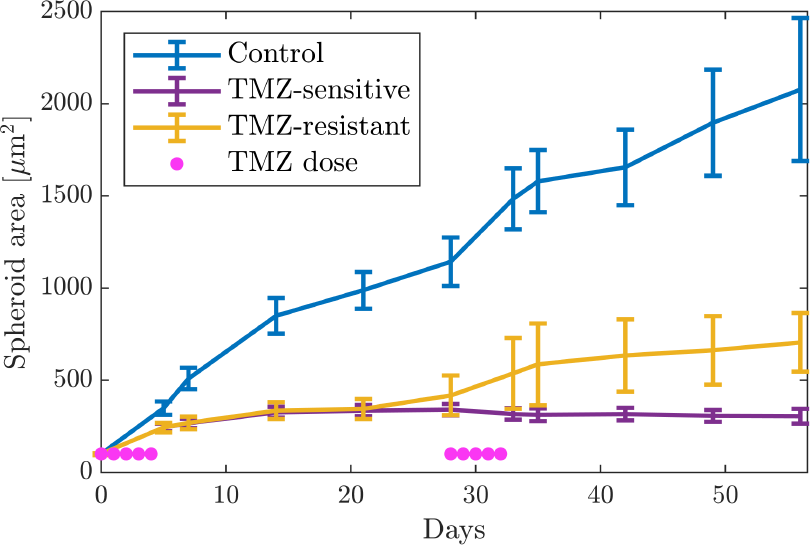
Experimental results. The curves show the temporal evolution of the spheroids area for each population, depicted as mean *±* standard deviation. The control group is composed of 143 spheroids. Regarding the spheroids treated with TMZ, there are 71 spheroids in the TMZ-sensitive group and 23 in the TMZ-resistant group.

### 2.2 Mathematical model

The guiding principle for developing a model describing spheroid evolution is the framework presented in [27] for modelling cell adaptation using internal variables. The main idea is to incorporate internal variables as macroscopic descriptors of the phenotypic state of cells. The evolution of these internal variables is driven by different fields associated with the specific microenvironment and cell epigenetic state. That is, the phenotype of cells affects their behaviour.

In this work, since we only have experimental data regarding the total size of the spheroids, we focus on homogeneous time-depending fields, neglecting spatial dependencies. The variables considered are then the number of alive and dead cells (obtained from the experimental data of the spheroid size assuming that the area is proportional to the number of cells and that cells do not disaggregate or change their size), the concentration of TMZ in the medium and an internal variable that takes into account the phenotypic changes caused by TMZ exposure.

Hereafter, the hypotheses on which the model is based are examined. According to experimental evidence and previous works [33, 34], we assume that only cells on the surface of the spheroid have enough space and resources to proliferate. This area of proliferative cells is often named as proliferative rim [20]. Because we do not have data on cell death, we assume, for simplicity, that the death rate is constant. Besides, it has been proved experimentally that spheroid proliferation slows down when cells are in contact with dead cells that secrete growth inhibition factors [33, 35]. Finally, we consider the possibility that some alive cells may be shed from the spheroid and that cells that have been dead for a certain period of time may dissolve.

With all the above, we define the equation governing the temporal evolution of the number of alive *N* (*t*) and dead *D*(*t*) cells forming the spheroid as:

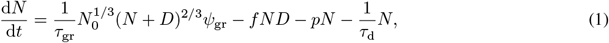

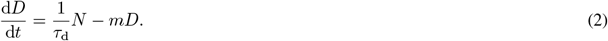

In Eq. (1), *τ*_gr_ is the characteristic time of cell proliferation, and *N*_0_ is the initial number of cells forming the spheroid, acting as a constant to standarise units. Cell proliferation is affected by the phenotypic state through function *ψ*_gr_, which will be detailed later. The second term in the RHS represents growth inhibition induced by cell death, regulated by the coefficient *f* . The third term represents the cells that are shed from the spheroid, regulated by the coefficient *p* and the last term is the death term, with *τ*_d_ the death characteristic time.

Regarding Eq. (2), the first term in the RHS represents the rate of formerly alive cells dying, while the second term represents the eventual disappearance (reabsorption) of dead cells through the coefficient *m*.

Next, we consider the evolution of the drug TMZ, the only substance that we consider relevant in the tumour microenvironment for this problem. The concentration of TMZ in the media (*T*) evolves according to:

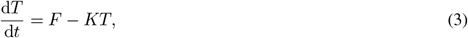

where *F* = *F* (*t*) indicates the dose administered at each time step *t* and *K* is the decay rate.

As commented in Section 2.1, in the in vitro experiments, a dose of *α* = 100 *μ*M is administered in days 𝒟 = {0, 1, 2, 3, 4, 27, 28, 29, 30, 31} . TMZ administration is modelled as an impulse since it is introduced instantaneously by changing the spheroid’s medium. Thus, *F* (*t*) is modelled with a Dirac Delta as:

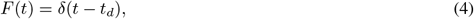

with *t*_*d*_ ∈ *𝒟* so that

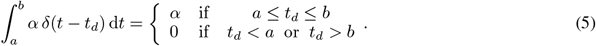

Next, we define the model hypotheses concerning the evolution of cell state through the internal variable. In this case, the state is only determined by the environmental stress caused by the epigenetic changes that TMZ causes in cells.

Cells undergo these changes when the concentration of TMZ in the surrounding medium is above a certain threshold. At the same time, cells adapt to their environment as they receive drug, and the more accumulated stress they have suffered, the more TMZ they are able to withstand without suffering additional stress. When cells accumulate a certain amount of epigenetic changes (that is, when the stress goes above a certain level), there is a cytostatic effect that causes the cells in the spheroid to stop proliferating. However, cells that develop resistance are able to repair those epigenetic changes [36, 37]. Last, we introduce the hypothesis that the cytostatic effect depends on the accumulated stress that cells have suffered, and not only on the current level, which could explain that, in the experiments, resistant cells do not grow as fast as control cells, even if the current stress level is not high.

With all this, we can write the equation for the evolution of the cell stress *S* as:

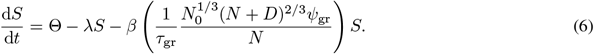

In Eq. (6) the first term models the acquisition of stress, which depends on the concentration of TMZ and is modelled as a Heaviside function regulated by the coefficient *κ*:

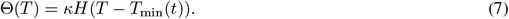

As previously commented, the TMZ threshold above which cells accumulate stress is variable and depends on the total amount of stress they have withstood. Thus:

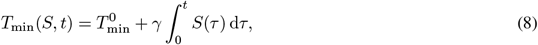

where *γ* is the adaptation coefficient, relating the increase in the threshold with the stress history. Besides, we assume that, initially, cells respond to any concentration of TMZ, so 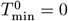.

In addition to cell adaptation, the cell’s ability to repair stress (the epigenetic changes suffered) is considered and consequently, a term of decay is included in Eq. (6), with *λ* the decay rate. Finally, the last term in Eq. (6) considers that epigenetic changes may be inherited. This may be seen as another way of repair, since when the changes are not inheritable, the population repairs itself through proliferation. Inheritance is regulated by the parameter *β*, which is bounded in [0, 1]. Indeed, when *β* = 0 there is no repair through inheritance and daughter cells are born with the same stress and epigenetic changes as their progenitors. However, when *β >* 0 cells do not pass on all their epigenetic changes to their offspring, which can be seen as another repair mechanism.

It still remains to define the function regulating the cytostatic effect in cells, that is, *ψ*_gr_ in Eq. (1). This function takes the form:

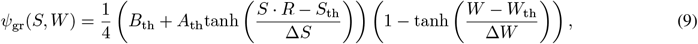

We model this effect at the population level, so we consider it as gradual rather than sharp, and thus we use hyperbolic tangent functions. The first hyperbolic tangent in Eq. (9) represents the cytostatic effect induced by the current level of stress. *S*_th_ is the characteristic stress threshold, marking the transition between normal and cytostatic growth, and Δ*S* is a spread parameter. *A*_th_ and *B*_th_ are defined to ensure that the value of the function goes to 1*/τ*_gr_ as *S* → 0 and to 1*/τ*_s_ as *S* →∞, being *τ*_s_ ≫ *τ*_gr_ the characteristic proliferation time when there is a cytostatic effect in the population ^1^ .

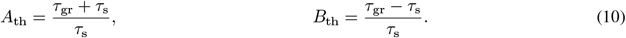

It has been highlighted in Section 2.1 that there is a certain delay between the administration of TMZ and the apparition of the cytostatic effect. To take this into account, we introduce a lag function *R*:

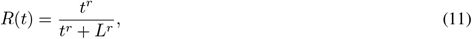

with *L* the lag time and *r* a coefficient determining the sharpness of the function. Finally, the second hyperbolic tangent models the effect of the accumulated stress *W* on proliferation (the fact that the cytostatic effect could depend on the history of stress):

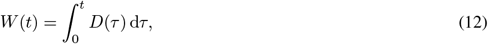

and *W*_th_ and Δ*W* are the threshold and spread parameter for this hyperbolic tangent respectively.

This model should be completed with suitable initial conditions, which, for the experiment described, are:

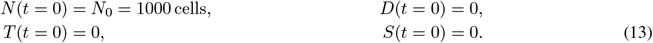

Table 1 shows a summary of the model parameters.

**Table 1:** Summary of model parameters. The whole set of parameters involved in the general model, together with their symbol and units.

| Parameter | Symbol | Units |
| --- | --- | --- |
| Growth characteristic time | $\tau_{gr}$ | h |
| Shedding rate | $p$ | $h^{-1}$ |
| Growth inhibition coefficient | $f$ | $cell^{-1}h^{-1}$ |
| Death characteristic time | $\tau_d$ | h |
| Dead cells disappearance rate | $m$ | $h^{-1}$ |
| TMZ decay rate | $K$ | $h^{-1}$ |
| Stress acquisition coefficient | $\kappa$ | $h^{-1}$ |
| Adaptation coefficient | $\gamma$ | $\mu M \cdot h^{-1}$ |
| Stress decay rate | $\lambda$ | $h^{-1}$ |
| Cytostatic growth characteristic time | $\tau_s$ | h |
| Lag time | $L$ | h |
| Lag sharpness coefficient | $r$ | - |
| Stress threshold | $S_{th}$ | - |
| Stress spread parameter | $\Delta S$ | - |
| Accumulated stress threshold | $W_{th}$ | h |
| Accumulated stress spread parameter | $\Delta W$ | h |
| Inheritance parameter | $\beta$ | - |

### 2.3 Fitting procedure and model selection

In the previous section, we have introduced a general model, including several phenomena corresponding to different assumptions and their corresponding parameters, which need to be calibrated with the experimental data. However, the actual underlying model is unknown and all these phenomena may not be necessary for accurately describing the data. Indeed, in many cases, models describing complex biological phenomena have too many parameters, leading to overly-complex models and overfitting, making it difficult to draw general conclusions [38, 29, 39]. For this reasons, it is paramount to follow a methodology for model selection. Hence, the objective of this section is twofold. On the one hand, we want to determine the most parsimonious model among a set of model candidates, obtained starting from the general (or full) model presented when neglecting some phenomena. On the other hand, we aim at finding the optimal value of the parameters that fit the experimental data presented in Section 2.1.

To select the most parsimonious model from a collection of different models, we use the Bayesian Information Criterion (BIC), since it is a consistent and unbiased estimator, taking into account both model complexity and model performance [40], and has been widely used for model selection in biological problems similar to the one in hands [28, 41, 29]. The BIC is expressed in terms of the chi-squared statistic (*χ*^2^) and a penalty term accounting for model complexity as:

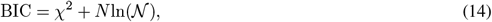

with *N* the number of free parameters to be estimated and *𝒩* the number of experimental data or observations. In our case, *𝒩* = 10 for each population, since there are 11 data points, but the first is used as initial condition.

Besides, we perform the model selection and parameter optimisation procedure in two stages. We divide the parameter set ***λ*** in two groups ***λ*** = (***λ***_1_, ***λ***_2_), depending on whether they are related to spheroid evolution (***λ***_1_) or to the response to treatment with TMZ (***λ***_2_). The TMZ decay rate parameter (*K* = 1.8 h) is not part of the optimisation since its value depends on the chemistry of the molecule and has been reported in the literature [42]. First, we optimise the parameters related to spheroid evolution (i.e., cell proliferation and death) using the data from the control spheroid population, since these spheroids are not treated and there are fewer coupled phenomena. This allows to calibrate the parameters avoiding that their effect is mixed with those of other phenomena. The parameters to be fitted in this stage are ***λ***_1_ = (*τ*_gr_, *τ*_d_, *f, p, m*), and the set of candidate models is obtained by considering all combinations by which we can remove several phenomena (except for growth and death, which are assumed to be always relevant):

- **Model 1a:** Complete model, considering all the parameters.
- **Model 1b:** Model in which we neglect the effect of dead cells that disappear from the spheroid (*m* = 0).
- **Model 1c:** Model in which we neglect the effect of alive cells shed away from the spheroid (*p* = 0).
- **Model 1d:** Model in which we neglect the effect of inhibition of growth due to dead cells (*f* = 0).
- **Model 1e:** Model in which we neglect the effect of dead cells that disappear from the spheroid and of alive cells shed away from the spheroid (*m* = *p* = 0).
- **Model 1f:** Model in which we neglect the effect of dead cells that disappear from the spheroid and the inhibition of growth due to dead cells (*m* = *f* = 0).
- **Model 1g:** Model in which we neglect the effect of alive cells shed away from the spheroid and the inhibition of growth due to dead cells (*p* = *f* = 0).
- **Model 1h:** Model in which we only consider proliferation and death (*m* = *p* = *f* = 0).

Second, once the best model from stage 1 has been determined, and considering the parameters obtained from the first stage as constant (with their value fixed to the optimal value obtained), we calibrate the parameters related to the response to TMZ, which are ***λ***_2_ = (*L, n, k, S*_th_, Δ*S, τ*_s_, *λ, γ, W*_th_, Δ*W, β*). For this purpose, we use the experimental data from the two populations of treated spheroids: sensitive and resistant. To allow for the differences in the behaviour between both groups, we assume that TMZ-sensitive spheroids are not able to reverse the drug’s effect, and hence we impose for this population *λ* = 0. Regarding the candidate models, we consider removing the phenomena related to the acquisition and repair of phenotypic changes: cytostatic effect due to accumulated stress, adaptation to TMZ and repair by heritage. We consider that the rest of phenomena are inherent to the experimental observations and/or reported in literature and, therefore, they cannot be removed. Consequently, the set of models in this stage is:

- **Model 2a:** Complete model, considering all the parameters.
- **Model 2b:** Model in which we assume that there is no possibility of repair by inheritance. That is, cells inherit the same state than their progenitor (*β* = 0).
- **Model 2c:** Model in which assume that the cytostatic effect is caused by the current phenotypic state (or stress) of cells, and not by the accumulated stress (equivalent to removing the second hyperbolic tangent in Eq. 9, and thus eliminating parameters *W*_th_, Δ*W*).
- **Model 2d:** Model in which we assume that cells do not adapt to TMZ. That is, the threshold of TMZ concentration above which cells undergo phenotypic changes is always the same (*γ* = 0).
- **Model 2e:** Model in which we assume that there is no possibility of repair by inheritance and that the cytostatic effect is not caused by accumulated stress (*β* = 0 and removal of parameters *W*_th_, Δ*W*).
- **Model 2f:** Model in which we assume that there is no possibility of repair by inheritance and that cells do not adapt to TMZ (*β* = *γ* = 0).
- **Model 2g:** Model in which we assume that cells do not adapt to TMZ and that the cytostatic effect is not caused by accumulated stress (*γ* = 0 and removal of parameters *W*_th_, Δ*W*).
- **Model 2h:** Model in which we only consider the base phenomena in the response to TMZ, that is, acquisition of phenotypic changes (with lag), spontaneous repair and cytostatic effect caused by stress. (*β* = *γ* = 0 and removal of parameters *W*_th_, Δ*W*).

Once the candidates are defined, for each stage the next step is to optimise the parameters for each candidate. The optimisation is carried out using nonlinear least squares with the lsqnonlin function in Matlab, in particular, using the Levenberg-Marquardt algorithm [43]. The objective function to minimise is the *χ*^2^ statistic, measuring the discrepancy between experimental data, represented by the mean *μ*_*i*_ and standard deviation *σ*_*i*_ corresponding to the observations at time *t*_*i*_, and simulated data *û*_*i*_ for each time *t*_*i*_ in which we have experimental data (*t*_*i*_ ∈ *𝒟*). *χ*^2^ is defined as:

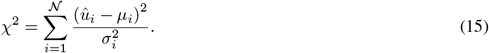

For the optimisation, all parameters have a lower bound of zero and no upper bound, except for *β*, which is bounded by its definition in the interval [0, 1]. When the optimisation finishes, the BIC is computed. Finally, the lowest BIC is selected among all candidates. A scheme of the optimisation and model selection procedure in two steps is represented in Figure 2.

**Figure 2:**
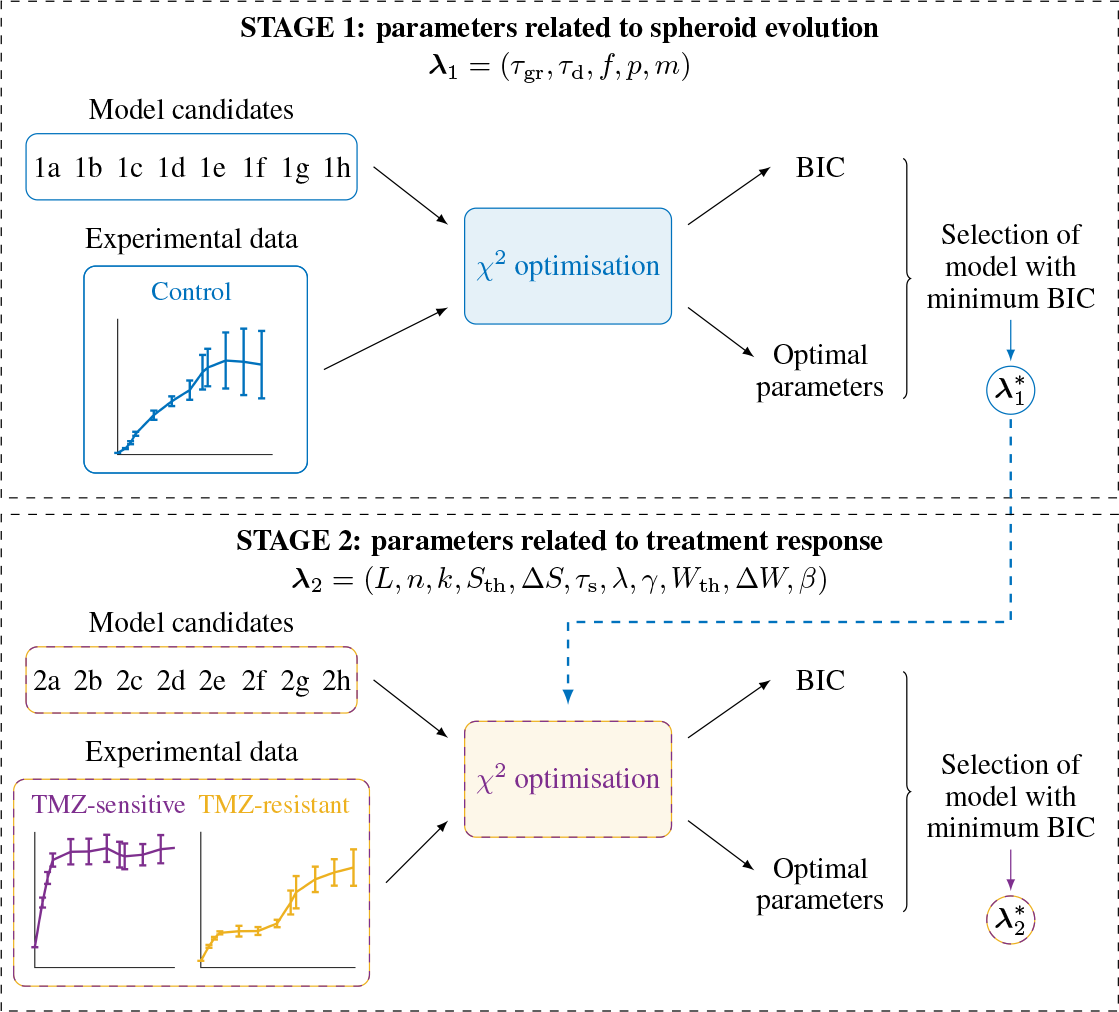
Schematic representation of the two-stages procedure followed for optimisation and model selection. As a result of this procedure, we obtain the set of optimal parameters 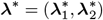.

### 2.4 Sensitivity analysis

After the best model is selected and the optimal value of the parameters is determined, we perform a sensitivity analysis to further inspect the model and how the uncertainty in the different parameters affects its output, understood as the total number of cells forming the spheroid.

After obtaining the optimal value for our parameters thanks to the available experimental data, we perform a local sensitivity analysis. The aim is to explore how the model output is affected by perturbations in the inputs around their nominal values (the fitted values) [44, 45]. We use a derivative-based method with one-at-a-time sampling, that is, only one parameter is perturbed at each time. The sensitivity index *S*_*i*_ for each parameter *λ*_*i*_ is then calculated with a partial derivative evaluated at the optimal value of the parameters ***λ***^*^:

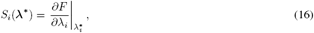

where *F* represents the model. This expression is usually numerically approximated as:

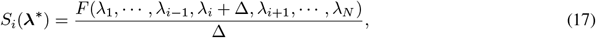

with Δ the magnitude of the perturbation.

## 3 Results

This section is dedicated to the presentation of the main findings obtained. In the first place, in Section 3.1 the results from the model selection procedure are showcased, finally selecting the model with the lowest BIC, and comparing experimental and simulated results obtained with the selected model. Then, in Section 3.2 the optimal parameter values are displayed together with contour plots of the *χ*^2^ statistic to examine how sensitive it is to perturbations of the parameters. Section 3.3 delves deeper into this aspect with a local sensitivity analysis around the optimal point in the parametric space. Finally, Section 3.4 addresses some biological questions that can arise and may be answered thanks to the mathematical model and methodology presented herein.

### 3.1 Model selection

Tables 2 and 3 show the results for the model selection procedure at stages 1 and 2 respectively. For the first stage, in which the phenomena and parameters not related to the treatment are taken into account, the model candidate with the lowest BIC value is 1e. We conclude that neither the shedding of alive cells (parameter *p*) away from the spheroid nor the decay of dead cells (parameter *m*) improve the fitting in a significant manner, so they will be neglected. Accordingly, in the final model, the evolution of untreated spheroids is characterised by growth, death and the inhibition of growth induced by dead cells.

**Table 2:** Summary of stage 1 of the optimisation and model selection procedure. The table summarises the different model candidates in stage 1, indicating which parameters are not considered in each candidate, together with its degrees of freedom (*ν*_1_) and the obtained metrics, the chi-squared statistic 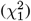 and the value of the Bayesian Information Criterion (BIC).

| Parameter | $\tau_{gr}$ | $\tau_d$ | $f$ | $p$ | $m$ | $\nu_1$ | $\chi_1^2$ | BIC |
| --- | --- | --- | --- | --- | --- | --- | --- | --- |
| Model 1a |  |  |  |  |  | 5 | 2.77 | 14.29 |
| Model 1b |  |  |  |  | 0 | 6 | 2.79 | 12.00 |
| Model 1c |  |  |  | 0 |  | 6 | 2.95 | 12.16 |
| Model 1d |  |  | 0 |  |  | 6 | 48.33 | 57.54 |
| Model 1e |  |  |  | 0 | 0 | 7 | 2.84 | 9.75 |
| Model 1f |  |  | 0 |  | 0 | 7 | 49.22 | 56.13 |
| Model 1g |  |  | 0 | 0 |  | 7 | 52.51 | 59.42 |
| Model 1h |  |  | 0 | 0 | 0 | 8 | 57.13 | 61.74 |

**Table 3:** Summary of stage 2 of the optimisation and model selection procedure. The table summarises the different model candidates in stage 2, indicating which parameters are not considered in each candidate, together with its degrees of freedom (*ν*_2_) and the obtained metrics, the chi-squared statistic 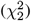 and the value of the Bayesian Information Criterion (BIC).

| Parameter | $L$ | $n$ | $\kappa$ | $S_{th}$ | $\Delta S$ | $\tau_s$ | $\lambda$ | $\gamma$ | $W_{th}$ | $\Delta W$ | $\beta$ | $\nu_2$ | $\chi_2^2$ | BIC |
| --- | --- | --- | --- | --- | --- | --- | --- | --- | --- | --- | --- | --- | --- | --- |
| Model 2a |  |  |  |  |  |  |  |  |  |  |  | 9 | 6.42 | 39.37 |
| Model 2b |  |  |  |  |  |  |  |  |  |  | 0 | 10 | 4.58 | 34.54 |
| Model 2c |  |  |  |  |  |  |  |  | 0 | 0 |  | 11 | 5.15 | 32.11 |
| Model 2d |  |  |  |  |  |  |  | 0 |  |  |  | 10 | 7.78 | 37.74 |
| Model 2e |  |  |  |  |  |  |  |  | 0 | 0 | 0 | 12 | 4.28 | 28.24 |
| Model 2f |  |  |  |  |  |  |  | 0 |  |  | 0 | 11 | 6.72 | 33.68 |
| Model 2g |  |  |  |  |  |  |  | 0 | 0 | 0 |  | 12 | 7.20 | 31.16 |
| Model 2h |  |  |  |  |  |  |  | 0 | 0 | 0 | 0 | 13 | 9.35 | 30.32 |

Similarly, for the second stage, the selected model is candidate 2e, which implies that cells inherit the same phenotypic state as their progenitors (*β* = 0), that is, there is no reversibility of the epigenetic changes through inheritance. Besides, the effect of the accumulated stress in proliferation (parameters *W*_th_ and Δ*W*) is also disregarded. In the final model, the cytostatic effect is caused only by the current (and not accumulated) phenotypic state.

Figure 3 shows the comparison between the experimental and simulated results, where it can be seen that we achieve a high level of agreement between both of them for the three populations, using a single set of parameters (except for the value of *λ* which, as explained before, was set to zero for the TMZ-sensitive population).

**Figure 3:**
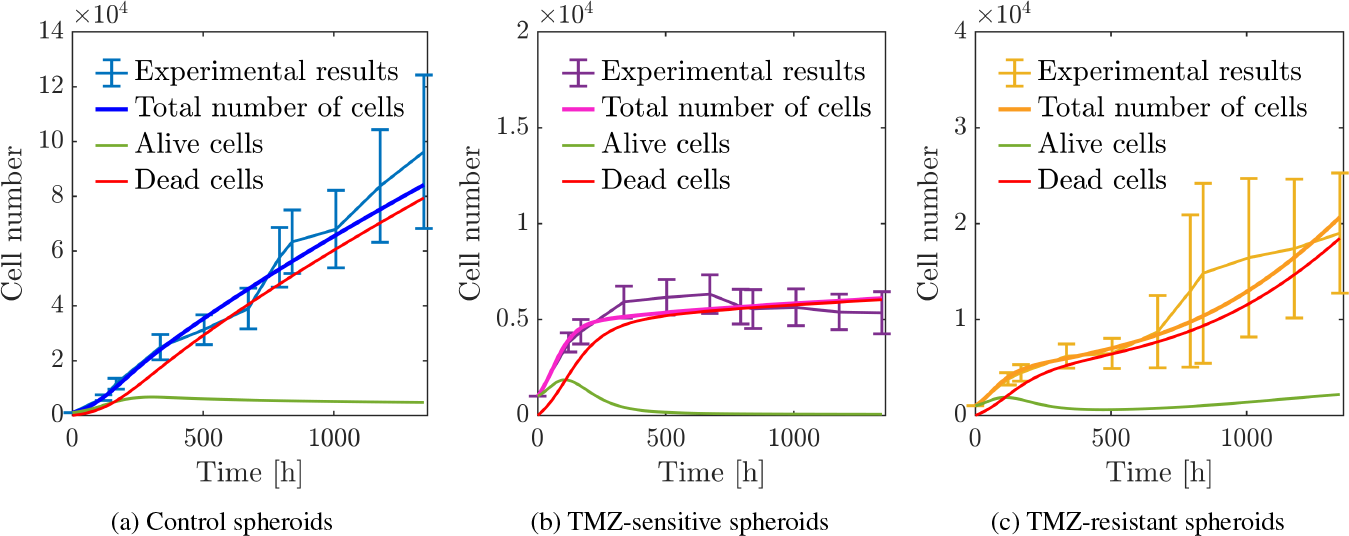
Comparison between experimental and simulated results. In each subfigure, the evolution of the number of alive (green line) and dead (red line) cells predicted by the model are represented, and the total number of cells is obtained as the sum of alive and dead cells.

To endorse the hypothesis that the model is able to accurately reproduce the experimental data, a hypothesis test was carried out. The null hypothesis (*H*_0_) is that the in silico data correctly describe the in vitro observations. To reject *H*_0_, it is required that 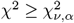, where 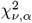 is the theoretical distribution of *χ*^2^ with *ν* degrees of freedom and significance level *α*, and *χ*^2^ as a weighted combination between 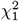 and 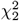:

In our case, *ν* = 𝒩 −*N* = *ν*_1_ + *ν*_2_ = 19. With this, we obtain a p-value of 0.99 and consequently there is not statistical evidence to reject *H*_0_.

### 3.2 Parameter fitting

One the final model is selected, and after following the optimisation procedure described in Section 2.3 to minimise the *χ*^2^ statistic, we obtain the optimal parameters, which are represented in Table 4.

**Table 4:** Fitted values of the model parameters. Value sof the parameters ***λ***^*^ fitted to the experimental data with the described optimisation procedure.

| Parameter | Fitted value | Units |
| --- | --- | --- |
| $\tau_{gr}$ | 42.8 | h |
| $f$ | $1.1 \times 10^{-6}$ | cell.$^{-1}$h$^{-1}$ |
| $\tau_d$ | 88.3 | h |
| $\kappa$ | $1.3 \times 10^{-3}$ | h$^{-1}$ |
| $\gamma$ | 1.0 | μMh$^{-1}$ |
| $\lambda$ | $8.6 \times 10^{-4}$ | h$^{-1}$ |
| $\tau_s$ | $5.8 \times 10^9$ | h |
| $L$ | 181.8 | h |
| $r$ | 1.7 | - |
| $S_{th}$ | 0.28 | - |
| $\Delta S$ | 0.40 | - |

Contour plots of the *χ*^2^ value are represented in Figure 4 (in 2D) and Figure 5 (in 3D) for the parameters related with the response to TMZ: the epigenetic changes acquisition rate *κ*, the adaptation rate *γ* and the parameters regulating the cytostatic effect in cells *S*_th_ and Δ*S*. The repair rate *λ* was not considered in this analysis since its value is imposed for the sensitive population. They are obtained sampling the parametric space and obtaining the value of *χ*^2^ for the different points. The contour plots are represented for three different probabilities (*p* = 0.5, *p* = 0.7 and *p* = 0.9)).

**Figure 4:**
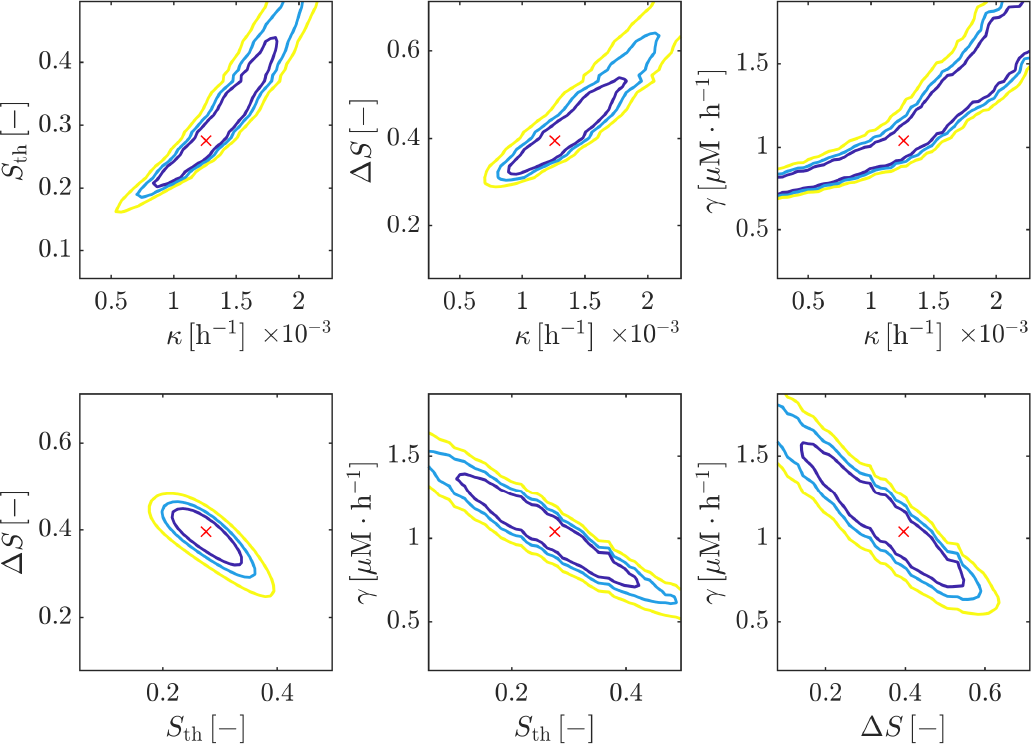
2D contour plots of the *χ*^2^ statistic. The contours are plotted for *p* = 0.5 (yellow), *p* = 0.7 (blue) and *p* = 0.9 (purple) for each pair of parameters in [*κ, γ, S*_th_, Δ*S*]. The red cross represents the optimal value of *χ*^2^.

**Figure 5:**
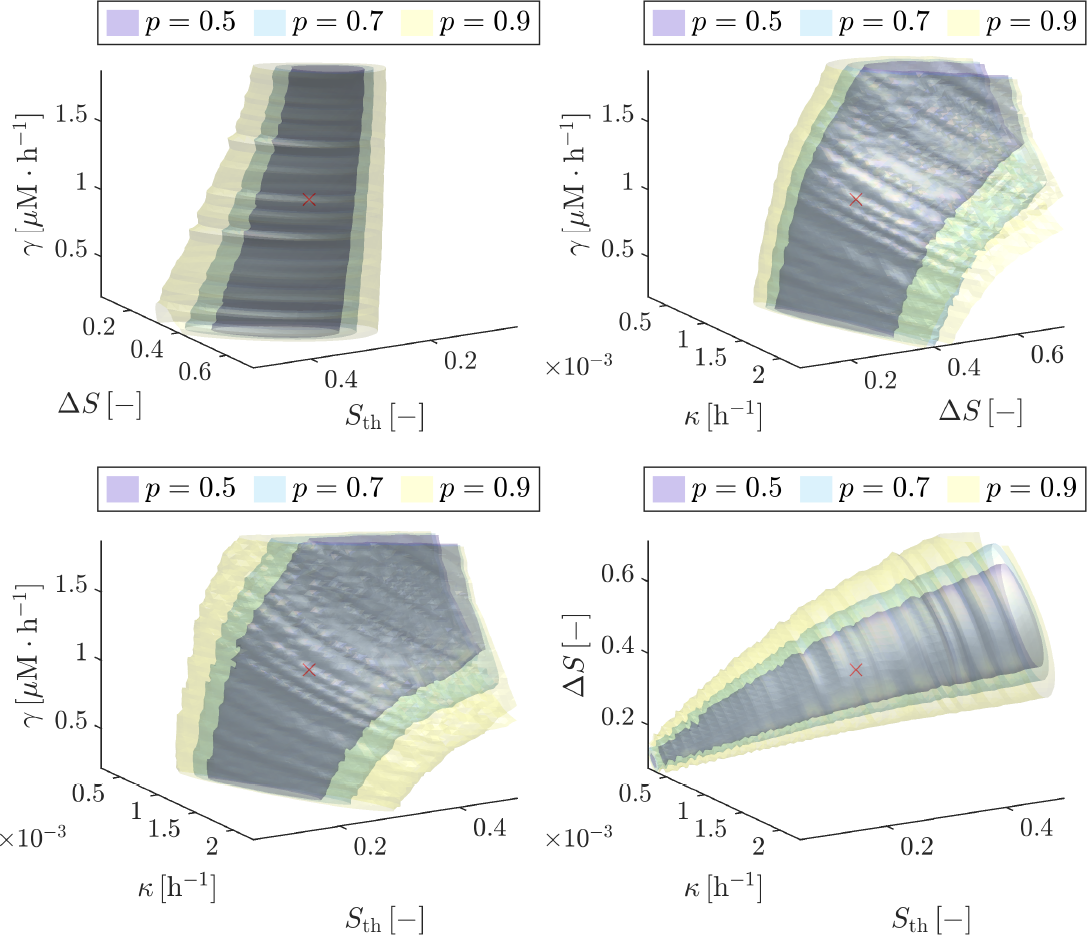
3D contour plots of the *χ*^2^ statistic. The contours are plotted for *p* = 0.5 (yellow), *p* = 0.7 (blue) and *p* = 0.9 (purple) for each parameter trio in [*κ, γ, S*_th_, Δ*S*]. The red cross represents the optimal value of *χ*^2^.

Figure 4 represents the contour curves for each combination of two parameters in [*κ, γ, S*_th_, Δ*S*], leaving the rest fixed at their fitted value (Table 4). As can be seen in the figure, the pair *S*_th_, Δ*S* has the lowest influence at the value of *χ*^2^ (always with the value of the rest of the parameters kept constant), while an increase in *κ* implies increasing in every other parameter to keep the same value of *χ*^2^. This is explained by the fact that an increase in the epigenetic changes acquisition rate leads to faster phenotypic changes and hence, to obtain the same results and mitigate this effect, we need either to increase the threshold for the cytostatic effect, or to increase the adaptation rate. On the other hand, an increment in the value of *γ* must be accompanied by a decrease in the parameters regulating the cytostatic effect, meaning that for higher adaptation rates (cells adapt faster to TMZ and thus are subject to less stress), this effect must occur for lower stress values.

In turn, Figure 5 represents three-dimensional contour curves for each parameter trio in [*κ, γ, S*_th_, Δ*S*]. This representation, although more difficult to interpret that the two-dimensional one, allows a deeper analysis of the interactions between three parameters at a time. For example, it can be seen that for [*γ, κ*, Δ*S*], Δ*S* has the lowest influence.

### 3.3 Sensitivity analysis

In this section we present the results of the local sensitivity analysis, in terms of the derivative-based sensitivity index (SI) defined in Section 2.4, computed with a perturbation of 10% of the optimal value of each parameter. In Figure 6 the normalised SI is represented for each of the parameters implied in the response to TMZ, for the sensitive (Figure 6a) and resistant (Figure 6b) spheroid populations, as well as the average of the indices of both populations (Figure 6c). The indices are similar for both populations, except of course for the index corresponding to *λ*, which has no effect in the sensitive population. Besides, the perturbation of *τ*_s_ has no effect either, given that its value is high enough to prevent spheroid growth in the considered time scale. The parameter whose perturbation has the higher impact is *κ*, which controls the level of the internal variable. It is also worth highlighting that, for the resistant group, the perturbation of the decay parameter *λ* has great influence in the results, since it controls how cells repair the epigenetic changes and eventually develop resistance to the drug. The effect and importance of this parameter will be further explored in Section 3.4.

**Figure 6:**
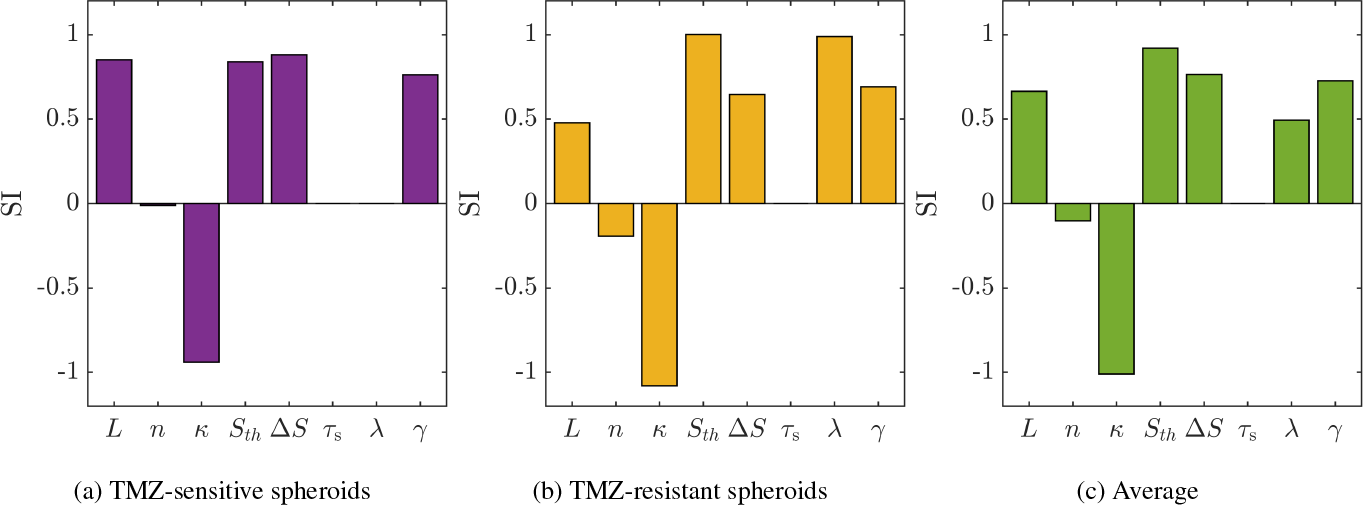
Sensitivity Index (SI) at the final simulation time. SI have been computed for each of the parameters involved in the response to TMZ. Subfigure (a): SI for the group TMZ-sensitive spheroids. Subfigure (b): SI for the group TMZ-resistant spheroids. Subfigure (c): Averaged SI for sensitive and resistant spheroids.

In order to get a more complete picture of the SIs, we evaluate the sensitivity of the complete temporal profile in Figure 7, where the SI of the evolution of the number of cells is represented.

**Figure 7:**
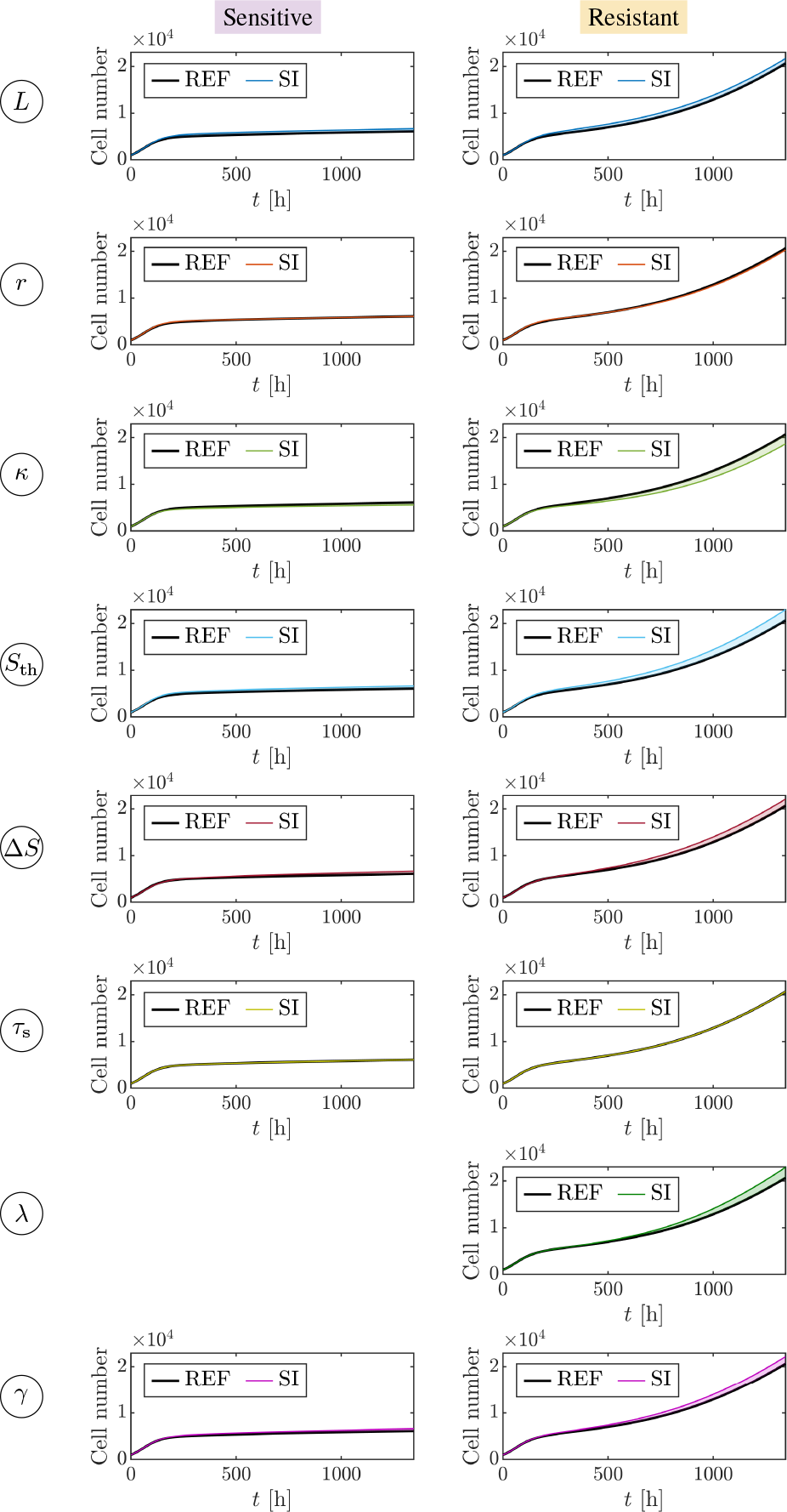
Sensitivity Index (SI) along time for each parameter. The SI have been computed for both sensitive (left column) and resistant (right column) spheroids and for each parameter (different rows) involved in the response to TMZ. The black line in each figure represents the simulation with the fitted parameters, taken as reference.

### 3.4 Biological insights from the model

This section is dedicated to exploring biological conclusions that can be drawn from the model. Two questions regarding the differences between sensitive and resistant populations are addressed, showing the capabilities of mathematical models such as the one presented in this work. First, we tackle the separation of treated spheroids in resistant and sensitive populations and then, we examine which mechanism is more likely to differentiate both populations.

#### 3.4.1 Is it possible to separate the treated populations in sensitive and resistant without prior information?

In the model selection and fitting procedure previously described, we started from experimental data which had been previously classified and separated in groups according to their response to TMZ and gene expression [32]. However, these techniques may not be available in every situarion and thus, in this section we want to classify treated spheroids in sensitive and resistant without using the prior available information.

For this purpose, we use the raw data, that is, the temporal evolution of the size of each individual spheroids (up to a total of 92 spheroids). The hypothesis, in line with the ones commented in previous sections, is that the populations can be separated using the value of the repair parameters [*β, λ*]. Hence, we first fit the value of the rest of the parameters involved with the response to TMZ using the mean size values of all spheroids. Once this done, we fit the value of the repair parameters for each individual spheroid, obtaining 92 pairs of values [*β, λ*].

To perform the clustering, we applied the k-means (*k* = 2) algorithm to the obtained bivariate parameter distribution, normalised with respect to the mean and the standard deviation of each parameter distribution obtained in the fitting (we call the normalised parameters 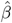 and 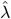). Based in our hypotheses, we assign the resistant class to the centroid with a higher value of 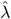 and 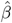. Performing different initialisations of the algorithm, we obtained between 1 and 2 incorrectly classified spheroids, if the separation proposed in [32] is assumed as correct.

The results for the best initialisation are summarised in Figure 8, together with the confusion matrix, with only one error of a resistant spheroid classified as sensitive. In this figure, the color of the marker represents the real group (purple for sensitive or yellow for resistant) to which each spheroid belongs, while the shape represents the assigned cluster (circle for sensitive and diamond for resistant). It can be seen that the spheroid which was wrongly classified (yellow circle) lies in the boundary between both clusters. Indeed, looking at the experimental profiles in subfigure (a), it becomes apparent that the spheroid that was incorrectly classified (whose evolution is highlighted with a red line) has the lowest final size among the resistant spheroids and could be taken for a sensitive one.

**Figure 8:**
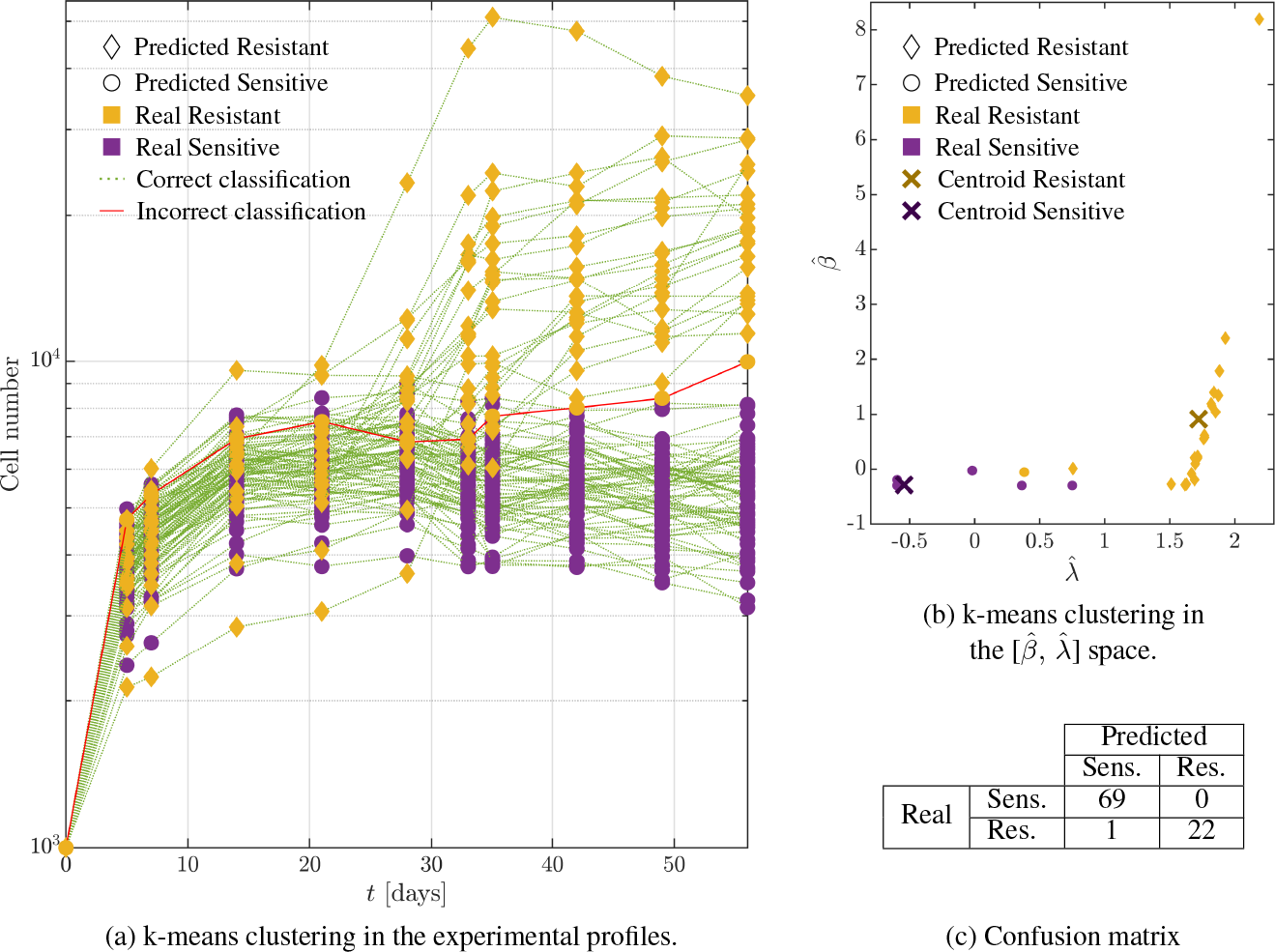
Results of the k-means clustering to classify the treated spheroids. Each experimental replicate was classified using the k-means algorithm depending on the normalised fitted value of the repair parameters 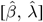. The parameters are normalised with the mean and standard deviation of the parametric sample. The color of the marker represents the real group to which each spheroid belongs, while the marker shapes represent the group assigned by k-means. Subfigure (a) shows the experimental profiles classified. Subfigure (b) shows the clustering in the parametric space 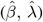. Subfigure (c) shows the confusion matrix.

To test whether the model provides a significant advantage for classification purposes with respect to the raw experimental data alone, we performed the clustering directly on the experimental data (the raw 92 temporal series). Trying again for 1 *×* 10^4^ different algorithm initialisations, we obtain between 7 (7.6%) and 22 (23.9%) incorrectly classified spheroids. Hence, we may conclude that using the proposed model and hypotheses could group the spheroids according to their response to treatment with TMZ in absence of gene expression data.

#### 3.4.2 Is repair spontaneous or driven by inheritance?

It is our hypothesis that sensitive and resistant spheroid populations differ in their ability to reverse or repair the epigenetic changes they acquire when exposed to TMZ, with TMZ-sensitive cells not being capable of any repair. In this regard, our framework includes two repair mechanisms, the spontaneous repair, mediated by parameter *λ* and the phenotypic state inheritance, mediated by parameter *β*.

The aim is to test which mechanism allows for better characterisation of both populations leading to a better fit of the experimental results. To do it, we consider different variations of the model, fit the results to each configuration and obtain the statistical descriptors of the goodness of fit, *χ*^2^ and BIC. We take as base model one in which there is no repair in any of the treated populations. Then, starting from this model, we study how the BIC is reduced introducing in the resistant populations each of the repair mechanisms described above, or both of them at the same time.

Table 5 summarises the results for the different hypotheses made. The hypothesis resulting in a lower BIC, that is, a bigger loss in the value of BIC with respect to the base value, is considering that the repair mechanism in resistant cells is spontaneous and that there is complete inheritance in both populations. This is coherent with the one of the predominant mechanisms behind TMZ-resistance, the over expression of MGMT [9]. That is, cells have pathways to repair epigenetic changes, such as the methylation induced by TMZ, not linked to division.

**Table 5:** Summary of BIC for different hypotheses concerning reversibility of the stress. The table summarises the different hypotheses, considering one, both or none of the possible repair mechanisms (inheritance and decay), together with the associated assumptions in terms of model parameters *β* and *λ*, the chi-squared statistic (*χ*^2^), the value of the Bayesian Information Criterion (BIC) and the BIC gain with respect to the base situation (first row), in which there is no possibility of repair.

| Hypothesis | Assumptions<br>TMZ-S | Assumptions<br>TMZ-R | $\chi^2$ | BIC | BIC loss |
| --- | --- | --- | --- | --- | --- |
| There is no repair in any form | $\beta = \lambda = 0$ | $\beta = \lambda = 0$ | 17.96 | 44.93 | - |
| Difference between sensitive and resistant is caused by the decay term and there is complete inheritance | $\beta = \lambda = 0$ | $\beta = 0$ | 4.28 | 28.24 | 16.69 |
| Difference between sensitive and resistant is caused by the inheritance term and there is no decay | $\beta = \lambda = 0$ | $\lambda = 0$ | 6.49 | 30.45 | 14.48 |
| Difference between sensitive and resistant is caused by the inheritance term and by the decay | $\beta = \lambda = 0$ | - | 8.81 | 35.76 | 9.17 |

These results are in line with those obtained from the k-means clustering in the previous section, where it could be seen (Figure 8, subfigure (b)) that the classification is made in terms of the value of *λ* and not of *β*.

## 4 Discussion

The work presented herein applies the methodology of model selection and calibration to a continuum model of cellular adaptation, with the aim of reproducing the acquisition of resistance to TMZ in GBM spheroids, as well as of broadening our understanding of the problem and how mathematical models can assist us in this task. The mathematical approach, first described in [27], tackles the modelling of cell phenotype as an internal variable which macroscopically represents the continuum epigenetic state of the cell. The general model has been particularised to our problem, considering as chemical species in the tumour microenvironment the concentration of the drug TMZ and an internal variable representing the epigenetic changes undergone as a result of drug exposition.

As opposed to other continuum approaches in literature [22, 23, 24, 25], which use different phenotypes for sensitive and resistant cells, the one here exposed is more biologically consistent, avoiding the assumption that cells instantaneously become resistant or sensitive [26]. There are some models already considering phenotype as a continuum for resistance modelling, but including phenotype as a structural dimension [46], instead of using internal variables. As discussed in [27], we believe that the presented approach offers straightforward implementation of the underlying biological hypotheses relating stimuli (the drug) and cell state, while having, in general, less parameters than multiphenotypic model, going in the direction of parsimony.

After the model selection and parameter fitting procedure followed, we were able to quantitatively reproduce the experimental results for the three populations (control, TMZ-senstitive and TMZ-resistant) with a single parameter set, as can be seen in Figure 3, achieving high levels of agreement. Only an extra assumption was required, in order to differentiate TMZ-sensitive and TMZ-resistant population, since this is a deterministic and not a stochastic model. For this reason, and following the biological hypothesis that TMZ-sensitive cells do not have the capability of reversing the methylation induced by TMZ, we impose that for this population the repair parameter *λ* is zero. It may be noted from Figure 3 that, by the final simulation time, the spheroids are mainly composed by necrotic cells. This is supported by experimental evidence holding that spheroids develop a necrotic core in the central area surrounded by a thin layer of proliferating cells (proliferating rim) [47]. The fitted value of the grow and death parameters is also in accordance with previous studies om in vitro culture of the U87 cell line [38, 48, 49].

One of the main hypothesis of the model is that cells adapt to the stimuli received, and that they are able to progressively withstand higher concentrations of TMZ without undergoing epigenetic changes [50]. This is modelled with a threshold *T*_min_ dependent on the accumulated exposure to the drug, which establishes the TMZ concentration above which epigenetic changes happen (and thus, the level of internal variable increases). Interestingly, if we take a look at the evolution of this threshold compared to the concentration of TMZ throughout the experiment (Figure 9), it becomes evident that for the whole second treatment cycle, the maximum drug concentration lies below the threshold. That is, the second cycle is not having effect on cell behaviour, suggesting that the same cells treated with only one cycle of TMZ between days 0 and 5 should exhibit a similar evolution to those treated with two cycles.

**Figure 9:**
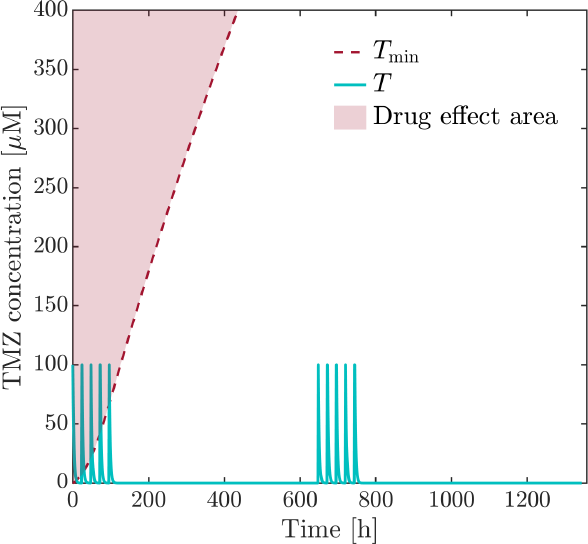
Temporal evolution of the TMZ concentration and the threshold for TMZ effect in cells. The blue line shows the evolution of the concentration of TMZ in the culture medium, while the red dashed line shows the threshold *T*_min_ above which cells undergo epigenetic changes due to drug exposure. The shaded area indicates where cells undergo these changes.

In fact, this was experimentally tested, and among treated spheroids there was also a group which developed resistance. In Figure 10 the evolution of the size of these spheroids is depicted, where it can be observed that the trend is indeed very similar, and that, as predicted by our model, only the first cycle is involved in the response of cells. This provides some validation for our approach, and exemplifies the utility of mathematical models such as the one here presented to propose new hypotheses and identifying the key mechanisms or variables in a complex process such as the acquisition of chemoresistance. This, in turn, helps reducing the number of experiments to be carried out in the laboratory, decreasing the associated costs, and pointing to the experiments which could potentially generate more knowledge. Of course, further validation would be required, to test if the model is able to generalise and correctly predict the spheroid evolution in different treatment schedules. Eventually, a fully validated model could act as a digital twin of the spheroids and used to test which would be the optimal treatment for achieving tumour remission.

**Figure 10:**
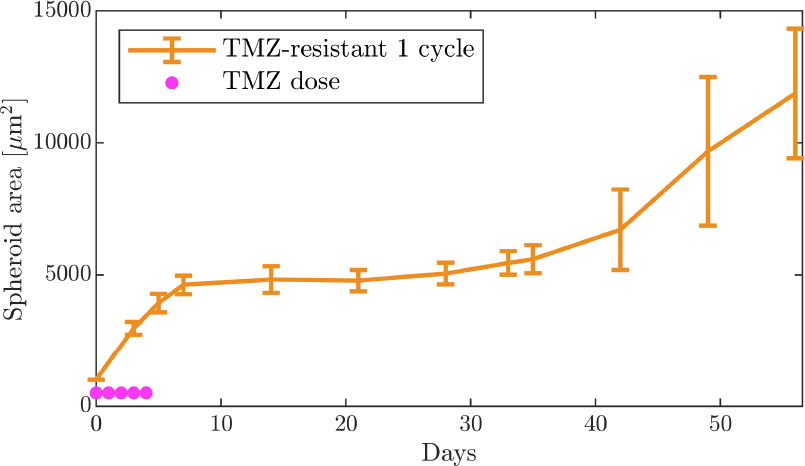
Experimental results of spheroids treated only with one cycle of TMZ. The temporal evolution of the resistant group of spheroids treated only with the first cycle of TMZ is shown. It can be seen how the trend is similar to those of resistant spheroids treated with two cycles of TMZ.

In addition to the example presented above, there are also other situations in which this model has proven to be useful. Indeed, in Section 3.4 we demonstrated that the model could be used to differentiate between sensitive and resistant spheroids with both high sensitivity and specificity (Figure 8), compared to the classification based in transcriptomics carried out in [32]. This is an example of how using the adequate mathematical techniques can be helpful to elucidate biological questions. A key idea here is that having a mathematical model allows reducing the dimensionality of the latent space in which the classification is carried out. Without the model, we move in a space of *N* dimensions, with *N* the number of experimental observations for each spheroid. However, our mathematical model and suitable hypotheses (the difference between both populations lies in the repair mechanisms) allows to reduce that space to a two dimensional space, which improves the classification outcome (reduction of the errors of at least 7-fold, as detailed in Section 3.4). This is an alternative to manifold learning techniques [51], as the latent space is built taking into account biological insights of the particular problem.

In Section 3.4 we also tackled a question about the most plausible repair mechanism (spontaneous repair or inheritance) to explain the differences between sensitive and resistant spheroids. The model selection procedure based on the BIC allows to resolve this question, being the model with the lowest BIC, in this case, the model with spontaneous repair in the resistant population, the most plausible one (Table 5). This result seems to be aligned with the previous one, in which the classification was made mainly in terms of the spontaneous repair parameter *λ*. However, we do not have experimental validation for this at the moment, being another example of how mathematical models can shade light in topics that are difficult to test in vitro [52, 53, 54]. However, even if we lack explicit validation, the finding is coherent with the over expression of MGMT often related to resistance, which is an spontaneous repair mechanism (in the sense that is internal to the cell and does not require external agents or cell division to occur).

In summary, the presented model allows quantitatively reproducing the process of resistance development in GBM spheroids subjected to treatment with TMZ, thanks to the definition of an internal variable modelling the stress acquired by cells when they are exposed to the drug. However, both the in vitro and in silico models used have some limitations. On the one hand, the in vitro model does not include all the typical GBM features, such as microvasculature or the interaction with the immune system [55, 56]. Besides, spheroids, albeit easy to handle, fail to recapitulate some clinical aspects such as irregular geometries, matrix remodelling or signalling pathways [57]. Hence, moving to patient specific frameworks with clinical data should be explored in the future, to bring the model closer to the clinical reality. On the other hand, the proposed model is homogeneous, since we only have data of the total spheroid size. This model could be extended to a partial differential equation model including spatial coordinates, given that we had data of the inner distributions of cells and TMZ inside the spheroid. Besides, the model could be extended to take into account other phenomena that are relevant in the process of resistance, such as the aforementioned angiogenesis or tumour-immune system interaction, as well as the inclusion of other chemical stimuli like oxygen. Indeed, hypoxia is a defining feature of GBM and is believed to play a role in the development of resistance [58, 59]. Despite these limitations, mathematical models such as the one presented herein, accompanied by an appropriate procedure for model selection and parameter fitting preventing overfitting, have been proven to be highly valuable for simulating cellular adaptation problems, that are key in cancer evolution [11]. Moreover, they are useful tools for addressing biological questions such as identifying populations or mechanisms.

## 5 Conclusion

Resistance to temozolomide is a major problem in the treatment of glioblastoma patients. At least 50 % of them do not respond to treatment and recurrence of the tumour is almost inevitable. Hence, overcoming temozolomide resistance is a major difficulty in the development of new treatment strategies.

In this work, we developed a mathematical model of cellular adaptation for the case of glioblastoma cells treated with temozolomide and their acquisition of resistance. The model is based on the concept of internal variables to model cell state, and it is fed with experimental data coming from spheroid culture of glioblastoma cells with two cycles of clinical treatment. Following a model selection methodology, we have been able of obtaining a final model able to accurately reproduce the evolution of the three populations of spheroids (control, resistant and sensitive), which contemplates the acquisition of epigenetic changes in response to drug exposure as well as the spontaneous repair of these changes, the adaptation to TMZ with stress and the cytostatic effect induced by phenotypic changes. The fitting results show that resistance is acquired with the first treatment cycle, and that the second cycle has no effect on cell behaviour, and indeed it was observed that treating cells only with one cycle yields similar results for in terms of evolution of resistant spheroids.

Indeed, we have been able to confirm the hypothesis of a difference between resistant and sensitive spheroids due to repair, since we are able to separate both populations based on the model’s repair parameters, highlighting the importance of mathematical models such as this for improving our understanding, isolating phenomena and checking hypotheses. The results suggest that the most relevant mechanism to explain the acquisition of resistance in the resistant spheroids is the so-called by us spontaneous repair, which models the internal pathways that cells have to revert epigenetic changes, being aligned with the existent literature.

In conclusion, the presented model and methodology, together with the selected examples and analyses, show the potential of the internal variables framework for modelling adaptation and phenotype for reproducing real data of one of the most crucial problems in cancer, the development of resistance. With further validation using different treatment schemes, the model could be used to perform in silico clinical trials that could save time and resources in the development of optimal treatments. Besides, we show the capabilities of the model for gaining insights into the underlying mechanisms, without using complex and expensive techniques.

## Acknowledgements

The authors gratefully acknowledge the financial support from the Spanish Ministry of Science and Innovation (MICINN), the State Research Agency (AEI), and FEDER, UE through the project PID2021-126051OB-C41.

## Footnotes

1 This parameter should tend to infinity for a complete cytostatic effect (no growth at all), but due to population effects, not all cells in the spheroid are equally sensitive, so this parameter is not assumed to be infinity but only sufficiently big.

## References

[1] Brian M Alexander and Timothy F Cloughesy. Adult glioblastoma. Journal of Clinical Oncology, 35(21):2402–2409, 2017.

[2] Ron Batash, Noam Asna, Pamela Schaffer, Nicole Francis, and Moshe Schaffer. Glioblastoma multiforme, diagnosis and treatment; recent literature review. Current medicinal chemistry, 24(27):3002–3009, 2017.

[3] Szymon Grochans, Anna Maria Cybulska, Donata Simińska, Jan Korbecki, Klaudyna Kojder, Dariusz Chlubek, and Irena Baranowska-Bosiacka. Epidemiology of glioblastoma multiforme–literature review. Cancers, 14(10):2412, 2022.

[4] Tijana Stanković, Teodora Ranđelović, Miodrag Dragoj, Sonja Stojković Burić, Luis Fernández, Ignacio Ochoa, Victor M Pérez-García, and Milica Pešić. In vitro biomimetic models for glioblastoma-a promising tool for drug response studies. Drug Resistance Updates, 55:100753, 2021.

[5] Shabierjiang Jiapaer, Takuya Furuta, Shingo Tanaka, Tomohiro Kitabayashi, and Mitsutoshi Nakada. Potential strategies overcoming the temozolomide resistance for glioblastoma. Neurologia medico-chirurgica, 58(10):405–421, 2018.

[6] Takahiro Oike, Yoshiyuki Suzuki, Ken-ichi Sugawara, Katsuyuki Shirai, Shin-ei Noda, Tomoaki Tamaki, Masaya Nagaishi, Hideaki Yokoo, Yoichi Nakazato, and Takashi Nakano. Radiotherapy plus concomitant adjuvant temozolomide for glioblastoma: Japanese mono-institutional results. PloS one, 8(11):e78943, 2013.

[7] Roger Stupp, Warren P Mason, Martin J Van Den Bent, Michael Weller, Barbara Fisher, Martin JB Taphoorn, Karl Belanger, Alba A Brandes, Christine Marosi, Ulrich Bogdahn, et al. Radiotherapy plus concomitant and adjuvant temozolomide for glioblastoma. New England journal of medicine, 352(10):987–996, 2005.

[8] Ian E McCutcheon and Mark C Preul. Historical perspective on surgery and survival with glioblastoma: how far have we come? World neurosurgery, 149:148–168, 2021.

[9] Maciej M Mrugala and Marc C Chamberlain. Mechanisms of disease: temozolomide and glioblastoma—look to the future. Nature clinical practice Oncology, 5(8):476–486, 2008.

[10] Shikhar Sharma, Theresa K Kelly, and Peter A Jones. Epigenetics in cancer. Carcinogenesis, 31(1):27–36, 2010.

[11] Douglas Hanahan. Hallmarks of cancer: new dimensions. Cancer discovery, 12(1):31–46, 2022.

[12] W Günther, E Pawlak, R Damasceno, H Arnold, and AJ Terzis. Temozolomide induces apoptosis and senescence in glioma cells cultured as multicellular spheroids. British journal of cancer, 88(3):463–469, 2003.

[13] Dorthe Aasland, Laura Götzinger, Laura Hauck, Nancy Berte, Jessica Meyer, Melanie Effenberger, Simon Schneider, Emelie E Reuber, Wynand P Roos, Maja T Tomicic, et al. Temozolomide induces senescence and repression of dna repair pathways in glioblastoma cells via activation of atr–chk1, p21, and nf-κb. Cancer research, 79(1):99–113, 2019.

[14] Sang Y Lee. Temozolomide resistance in glioblastoma multiforme. Genes & diseases, 3(3):198–210, 2016.

[15] Andrew Oliveira Silva, Eloisa Dalsin, Giovana Ravizzoni Onzi, Eduardo Cremonese Filippi-Chiela, and Guido Lenz. The regrowth kinetic of the surviving population is independent of acute and chronic responses to temozolomide in glioblastoma cell lines. Experimental Cell Research, 348(2):177–183, 2016.

[16] Khaled Messaoudi, Anne Clavreul, and Frédéric Lagarce. Toward an effective strategy in glioblastoma treatment. part i: resistance mechanisms and strategies to overcome resistance of glioblastoma to temozolomide. Drug discovery today, 20(7):899–905, 2015.

[17] Manendra Singh Tomar, Ashok Kumar, Chhitij Srivastava, and Ashutosh Shrivastava. Elucidating the mechanisms of temozolomide resistance in gliomas and the strategies to overcome the resistance. Biochimica et Biophysica Acta (BBA)-Reviews on Cancer, 1876(2):188616, 2021.

[18] Neha Singh, Alexandra Miner, Lauren Hennis, and Sandeep Mittal. Mechanisms of temozolomide resistance in glioblastoma-a comprehensive review. Cancer drug resistance, 4(1):17, 2021.

[19] Ana S Nunes, Andreia S Barros, Elisabete C Costa, André F Moreira, and Ilídio J Correia. 3d tumor spheroids as in vitro models to mimic in vivo human solid tumors resistance to therapeutic drugs. Biotechnology and bioengineering, 116(1):206–226, 2019.

[20] Sritama Nath and Gayathri R Devi. Three-dimensional culture systems in cancer research: Focus on tumor spheroid model. Pharmacology & therapeutics, 163:94–108, 2016.

[21] Teodora Ranđelović, Alodia Lacueva-Aparicio, Violeta Marques Larraz, Marina Perez-Aliacar, Isabel Marquina, Rebeca Sanz Pamplona, and Ignacio Ochoa. Heterogeneous response of glioblastoma spheroids to temozolomide treatment – chemoresistance development. Manuscript submitted for publication, 2023.

[22] Sharon S Hori, Ling Tong, Srividya Swaminathan, Mariola Liebersbach, Jingjing Wang, Sanjiv S Gambhir, and Dean W Felsher. A mathematical model of tumor regression and recurrence after therapeutic oncogene inactivation. Scientific reports, 11(1):1–14, 2021.

[23] Edouard Ollier, Pauline Mazzocco, Damien Ricard, Gentian Kaloshi, Ahmed Idbaih, Agusti Alentorn, Dimitri Psimaras, Jérôme Honnorat, Jean-Yves Delattre, Emmanuel Grenier, et al. Analysis of temozolomide resistance in low-grade gliomas using a mechanistic mathematical model. Fundamental & clinical pharmacology, 31(3):347–358, 2017.

[24] James M Greene, Jana L Gevertz, and Eduardo D Sontag. Mathematical approach to differentiate spontaneous and induced evolution to drug resistance during cancer treatment. JCO clinical cancer informatics, 3:1–20, 2019.

[25] Luis E Ayala-Hernández, Armando Gallegos, Philippe Schucht, Michael Murek, Luis Pérez-Romasanta, Juan Belmonte-Beitia, and Víctor M Pérez-García. Optimal combinations of chemotherapy and radiotherapy in low-grade gliomas: a mathematical approach. Journal of personalized medicine, 11(10):1036, 2021.

[26] Arran Hodgkinson, Laurent Le Cam, Dumitru Trucu, and Ovidiu Radulescu. Spatio-genetic and phenotypic modelling elucidates resistance and re-sensitisation to treatment in heterogeneous melanoma. Journal of theoretical biology, 466:84–105, 2019.

[27] Marina Pérez-Aliacar, Jacobo Ayensa-Jiménez, and Manuel Doblaré. Modelling cell adaptation using internal variables: Accounting for cell plasticity in continuum mathematical biology. Computers in Biology and Medicine, 164:107291, 2023.

[28] Anna Claudia M Resende, Ernesto ABF Lima, Regina C Almeida, Matthew T McKenna, and Thomas E Yankeelov. Model selection for assessing the effects of doxorubicin on triple-negative breast cancer cell lines. Journal of Mathematical Biology, 85(6-7):65, 2022.

[29] EABF Lima, JT Oden, D. Hormuth, TE Yankeelov, and RC Almeida. Selection, calibration, and validation of models of tumor growth. Mathematical Models and Methods in Applied Sciences, 26(12):2341–2368, 2016.

[30] Sandrine Ostermann, Chantal Csajka, Thierry Buclin, Serge Leyvraz, Ferdy Lejeune, Laurent A Decosterd, and Roger Stupp. Plasma and cerebrospinal fluid population pharmacokinetics of temozolomide in malignant glioma patients. Clinical cancer research, 10(11):3728–3736, 2004.

[31] David Lacalle, Héctor Alfonso Castro-Abril, Teodora Randelovic, César Domínguez, Jónathan Heras, Eloy Mata, Gadea Mata, Yolanda Méndez, Vico Pascual, and Ignacio Ochoa. Spheroidj: An open-source set of tools for spheroid segmentation. Computer Methods and Programs in Biomedicine, 200:105837, 2021.

[32] Teodora Randelovic. The Role of Microenviroment in Glioblastoma Progression and Resistance Development. Universidad de Zaragoza. Phd thesis, University of Zaragoza, 2022. Available at https://zaguan.unizar.es/record/118607.

[33] James P Freyer. Role of necrosis in regulating the growth saturation of multicellular spheroids. Cancer research, 48(9):2432–2439, 1988.

[34] Judah Folkman and Mark Hochberg. Self-regulation of growth in three dimensions. The Journal of experimental medicine, 138(4):745–753, 1973.

[35] Dorothy I Wallace and Xinyue Guo. Properties of tumor spheroid growth exhibited by simple mathematical models. Frontiers in oncology, 3:51, 2013.

[36] Qiong Wu, Anders E Berglund, and Arnold B Etame. The impact of epigenetic modifications on adaptive resistance evolution in glioblastoma. International journal of molecular sciences, 22(15):8324, 2021.

[37] Jihong Zhang, Malcolm FG Stevens, and Tracey D Bradshaw. Temozolomide: mechanisms of action, repair and resistance. Current molecular pharmacology, 5(1):102–114, 2012.

[38] Jacobo Ayensa-Jiménez, Marina Pérez-Aliacar, Teodora Randelovic, Sara Oliván, Luis Fernández, José Antonio Sanz-Herrera, Ignacio Ochoa, Mohamed H Doweidar, and Manuel Doblaré. Mathematical formulation and parametric analysis of in vitro cell models in microfluidic devices: application to different stages of glioblastoma evolution. Scientific Reports, 10(1):21193, 2020.

[39] Daniel J VandenHeuvel, Christopher Drovandi, and Matthew J Simpson. Computationally efficient mechanism discovery for cell invasion with uncertainty quantification. PLOS Computational Biology, 18(11):e1010599, 2022.

[40] Andrew A Neath and Joseph E Cavanaugh. The bayesian information criterion: background, derivation, and applications. Wiley Interdisciplinary Reviews: Computational Statistics, 4(2):199–203, 2012.

[41] Santiago D Cárdenas, Constance J Reznik, Ruchira Ranaweera, Feifei Song, Christine H Chung, Elana J Fertig, and Jana L Gevertz. Model-informed experimental design recommendations for distinguishing intrinsic and acquired targeted therapeutic resistance in head and neck cancer. npj Systems Biology and Applications, 8(1):32, 2022.

[42] JR Wesolowski, P Rajdev, and SK Mukherji. Temozolomide (temodar). American journal of neuroradiology, 31(8):1383–1384, 2010.

[43] Jorge J Moré. The levenberg-marquardt algorithm: implementation and theory. In Numerical analysis: proceedings of the biennial Conference held at Dundee, June 28–July 1, 1977, pages 105–116. Springer, 2006.

[44] Emanuele Borgonovo. Sensitivity analysis of model output with input constraints: A generalized rationale for local methods. Risk Analysis: An International Journal, 28(3):667–680, 2008.

[45] Francesca Pianosi, Keith Beven, Jim Freer, Jim W Hall, Jonathan Rougier, David B Stephenson, and Thorsten Wagener. Sensitivity analysis of environmental models: A systematic review with practical workflow. Environmental Modelling & Software, 79:214–232, 2016.

[46] Giulia L Celora, Helen M Byrne, and PG Kevrekidis. Spatio-temporal modelling of phenotypic heterogeneity in tumour tissues and its impact on radiotherapy treatment. Journal of Theoretical Biology, 556:111248, 2023.

[47] HS Bell, IR Whittle, M Walker, HA Leaver, and SB Wharton. The development of necrosis and apoptosis in glioma: experimental findings using spheroid culture systems. Neuropathology and applied neurobiology, 27(4):291–304, 2001.

[48] Ana Carrasco-Mantis, Teodora Randelovic, Héctor Castro-Abril, Ignacio Ochoa, Manuel Doblaré, and José A Sanz-Herrera. A mechanobiological model for tumor spheroid evolution with application to glioblastoma: A continuum multiphysics approach. Computers in Biology and Medicine, 159:106897, 2023.

[49] Shi-cang Yu, Yi-fang Ping, Liang Yi, Zhi-hua Zhou, Jian-hong Chen, Xiao-hong Yao, Lei Gao, Ji Ming Wang, and Xiu-wu Bian. Isolation and characterization of cancer stem cells from a human glioblastoma cell line u87. Cancer letters, 265(1):124–134, 2008.

[50] Chia-Hung Chien, Wei-Ting Hsueh, Jian-Ying Chuang, and Kwang-Yu Chang. Dissecting the mechanism of temozolomide resistance and its association with the regulatory roles of intracellular reactive oxygen species in glioblastoma. Journal of biomedical science, 28(1):1–10, 2021.

[51] Alan Julian Izenman. Introduction to manifold learning. Wiley Interdisciplinary Reviews: Computational Statistics, 4(5):439–446, 2012.

[52] Irene Lacal and Rossella Ventura. Epigenetic inheritance: concepts, mechanisms and perspectives. Frontiers in molecular neuroscience, 11:292, 2018.

[53] Luciane T Kagohara, Genevieve L Stein-O’Brien, Dylan Kelley, Emily Flam, Heather C Wick, Ludmila V Danilova, Hariharan Easwaran, Alexander V Favorov, Jiang Qian, Daria A Gaykalova, et al. Epigenetic regulation of gene expression in cancer: techniques, resources and analysis. Briefings in functional genomics, 17(1):49–63, 2018.

[54] Nicholas Redshaw, Jim F Huggett, Martin S Taylor, Carole A Foy, and Alison S Devonshire. Quantification of epigenetic biomarkers: an evaluation of established and emerging methods for dna methylation analysis. BMC genomics, 15(1):1–14, 2014.

[55] Dimitrios Strepkos, Mariam Markouli, Alexia Klonou, Christina Piperi, and Athanasios G Papavassiliou. Insights in the immunobiology of glioblastoma. Journal of Molecular Medicine, 98:1–10, 2020.

[56] Caterina Brighi, Simon Puttick, Stephen Rose, and Andrew K Whittaker. The potential for remodelling the tumour vasculature in glioblastoma. Advanced Drug Delivery Reviews, 136:49–61, 2018.

[57] Marzenna Wiranowska and Mumtaz V Rojiani. Extracellular matrix microenvironment in glioma progression. Glioma–exploring its biology and practical relevance, pages 257–284, 2011.

[58] Ahmed Musah-Eroje and Sue Watson. A novel 3d in vitro model of glioblastoma reveals resistance to temozolomide which was potentiated by hypoxia. Journal of Neuro-oncology, 142:231–240, 2019.

[59] Chii-Wen Chou, Chi-Chung Wang, Chung-Pu Wu, Yu-Jung Lin, Yu-Chun Lee, Ya-Wen Cheng, and Chia-Hung Hsieh. Tumor cycling hypoxia induces chemoresistance in glioblastoma multiforme by upregulating the expression and function of abcb1. Neuro-oncology, 14(10):1227–1238, 2012.

